# Chromosomal rearrangements preserve adaptive divergence in ecological speciation

**DOI:** 10.1101/2021.08.20.457158

**Authors:** Craig E. Jackson, Sen Xu, Zhiqiang Ye, Michael E. Pfrender, Michael Lynch, John K. Colbourne, Joseph R. Shaw

## Abstract

Despite increasing empirical evidence that chromosomal rearrangements may play an important role in adaptive divergence and speciation, the degree to which divergent genomic regions are associated with chromosomal rearrangements remains unclear. In this study, we provide the first whole-genome analyses of ecological speciation and chromosomal evolution in a *Daphnia* species complex, using chromosome-scale assemblies and natural-population sequencing of the recently diverged species pair, *Daphnia pulicaria* and *Daphnia pulex*, which occupy distinct yet overlapping habitats in North America, and the outgroup species *Daphnia obtusa*. Our results describe a mixed mode of geographic divergence (isolation with secondary contact) resulting in a heterogeneous landscape of genomic divergence. Large contiguous “continents of divergence” encompass over one third of the genome (36%) and contain nearly all the fixed differences (94%) between the species, while the background genome has been homogenized. Chromosomal rearrangements between species, including inversions and intrachromosomal translocations, are associated with the continents of divergence and capture multiple adaptive alleles in genes and pathways thought to contribute to the species’ phenotypic differences.

## Introduction

Chromosomal rearrangements, especially inversions, are of considerable interest in understanding adaptation and speciation because they form barriers to gene flow between related species^1–4^. Chromosomal rearrangements can suppress recombination in chromosomal heterozygotes^5^, thereby reducing introgression across rearranged regions and protecting favorable genotypic combinations within the rearranged regions from being broken by recombination^6–8^. Indeed, recent theory and simulations demonstrate how inversions harboring locally adapted alleles can be selectively favored over collinear arrangements when populations are exchanging migrants or hybridizing^9^. Consequently, inversions rise to high frequency and populations diverge. In addition, such adaptive spread of inversions can be further facilitated when inversions arise in allopatry and are maintained at low frequency, but subsequently rise to high frequency when gene flow ensues upon secondary contact^10^. Despite the increasing empirical evidence that rearrangements are important role for adaptive divergence^11–14^, the degree to which divergent genomic regions are associated with chromosomal rearrangements remains unclear across the tree of life.

Speciation with gene flow is expected to generate heterogeneous patterns of divergence along the genome, where divergent selection acts on a few genes and linked loci while gene flow homogenizes the remainder of the genome^15,16^. Characterizing the genome-wide pattern of divergence in young species pairs has offered insights into multiple aspects of the genomic architecture of speciation, including the arrangement, physical scale, function, and number of highly differentiated regions in the genome^17–19^. Highly differentiated regions in hybridizing sympatric species can exist as only a handful of narrow peaks, such as in the threespine sticklebacks^20^, *Ficedula* flycatchers^21^, and *Corvus* crows^22^. These patterns of genome divergence gave rise to the metaphor “genomic islands of divergence” embedded in a sea of baseline genetic divergence^23^. Alternatively, widespread divergence and selection throughout the genome can exist, such as in the *Rhagoletis* fruit flies^24^ and *Menidia* silversides^25^, displaying “continents” of multiple differentiated loci in linkage disequilibrium. Furthermore, recent advances in demographic modeling have provided insight into the evolutionary history that shapes these genomic patterns of divergence during speciation, including the variation in the rate of gene flow across the genome^26–28^.

*Daphnia pulex* was the first crustacean for which a reference genome was assembled ^29^, and the *D. pulex* species complex includes at least 10 distinct lineages distributed globally with an evolutionary history of hybridization and introgression^30–32^. *Daphnia* genome evolution is also rich in structural variation, characterized by a high and historically steady rate of gene duplication and gene-family expansion and diversification^29^. Mutation accumulation experiments also indicate that *D. pulex* has high mutational rates of complex, large-scale duplications and deletions^33^. Therefore, the *D. pulex* species complex offers a promising system for studying speciation with gene flow and the role of chromosomal rearrangements in speciation.

This study focuses on the cyclically parthenogenetic (CP) species pair, *D. pulicaria* and *D. pulex*, where both prezygotic and postzygotic reproductive incompatibilities have started to arise^34,35^, and strong genetic evidence exists for divergence with gene flow^36^. *Daphnia pulicaria* and *D. pulex* are morphologically nearly indistinguishable, but occupy distinct, overlapping habitats in North America, with *D. pulicaria* inhabiting stratified permanent lakes while *D. pulex* mostly live in ephemeral fishless ponds^37^. Moreover, these two species show clear phenotypic differences, suggesting local adaptation and divergent selection in their distinct habitats. For example, *D. pulex* grows faster, reproduces at an earlier age, and goes through sexual reproduction via resting eggs before ponds dry up in summer, whereas *D. pulicaria* have a longer life span and can persist in lakes largely without sex for extended time^35,38–41^. Inhabiting stratified lakes with possible fish predation, *D. pulicaria* also exhibit the anti-predator behavior diel vertical migration through negative phototaxis, migrating to deeper, darker waters during the day and returning to surface waters at night to feed^42–44^.

Here we provide comprehensive whole-genome analyses of divergence and speciation in a *Daphnia* species complex. Using an integrative chromosome-scale metapopulation genomics approach including population genetic statistics, demographic modeling, comparative genome synteny, and genome visualization, we reconstruct the evolutionary history of ecological speciation, highlight the complete genomic landscape of divergence, and uncover an important role of chromosomal rearrangements in preserving adaptive divergence.

## Results

### Chromosome-scale genome assembly of *Daphnia* genomes

In order to investigate the complete and contiguous genomic landscape of divergence, we generated a chromosome-scale assembly of *D. pulicaria* by combining PacBio SMRT, Illumina mate-pair, Dovetail Chicago, and Dovetail Hi-C sequencing (Supplemental Table 1). We also generated an improved *D. pulex* assembly using PacBio sequencing, which is more contiguous than the previously released *D. pulex* Illumina short read assembly^45^. In addition, we generated a chromosome-scale assembly of *D. obtusa*, a closely related outgroup species to *D. pulicaria* and *D. pulex*. These *Daphnia* genomes are known to consist of 12 chromosomes (2n = 24)^29,46^. Dovetail Hi-C sequencing resolved all 12 chromosome-scale scaffolds in *D. pulicaria*, ranging from 8 Mb to 21 Mb in length and accounting for 92% of the total assembly. Using mRNA transcript expression, protein domain signatures, and protein orthology, we predicted and functionally annotated 27,276 protein-coding genes in the *D. pulicaria* genome. Throughout this study, the 12 *D. pulicaria* chromosome-scale scaffolds are used as the basis to examine the genomic landscape of divergence and synteny between *D. pulicaria* and *D. pulex*, while using *D. obtusa* as an outgroup for our analyses.

### Differential introgression during secondary contact shapes the genomic landscape of divergence

We used whole-genome sequences of geographically widely distributed *D. pulicaria* (n=14) and *D. pulex* (n=12) isolates from North American populations to investigate the genomic landscape of divergence between the species (Supplementary Fig. 1, Supplementary Table 2). These sequences were mapped to the *D. pulicaria* assembly in order to investigate within- and among-population genomics across the 12 chromosomes. Population structure and species divergence were analyzed using 2,049,444 high quality bi-allelic nuclear SNPs (on average ~124 SNPs per 10 kb) with minor-allele frequencies > 0.05. In order to polarize alleles as ancestral vs. derived, we used whole-genome sequences from *D. obtusa* as an outgroup to the *D. pulex* species complex. A maximum-likelihood unrooted phylogeny based on these nuclear SNPs clearly showed that *D. pulex* and *D. pulicaria* formed their own distinct clades, with within-species relatedness largely corresponding to geographic proximity (Supplementary Fig. 1). Principal component analysis substantiated the species-specific clusters of *D. pulex* and *D. pulicaria*, with the primary component explaining 31.1% of the total genetic variance (Supplementary Fig. 1).

To understand the evolutionary dynamics of speciation with gene flow in this species pair, we inferred the historical demography of *D. pulex* and *D. pulicaria* using a composite likelihood approach^47^. Seven candidate demographic models were fitted to the genome-wide joint allele-frequency spectrum (Fig. 1a-b), including scenarios of strict isolation (*SI*), isolation-with-migration (*IM*), ancient migration (*AM*), and secondary contact (*SC*). The observed allele-frequency spectrum was not satisfactorily reproduced by these standard demographic models that assume the same gene flow parameters among all loci. To incorporate heterogeneous gene flow, we included the *IM2m, AM2m*, and *SC2m* divergence models that account for variation in gene flow across the genome using two categories of migration rates^26^. Notably, the *SC2m* model, in which gene flow occurred following an allopatric phase (Fig. 1c-d), produced the best fit compared with all other alternative models (Supplementary Table 3).

**Figure 1.**
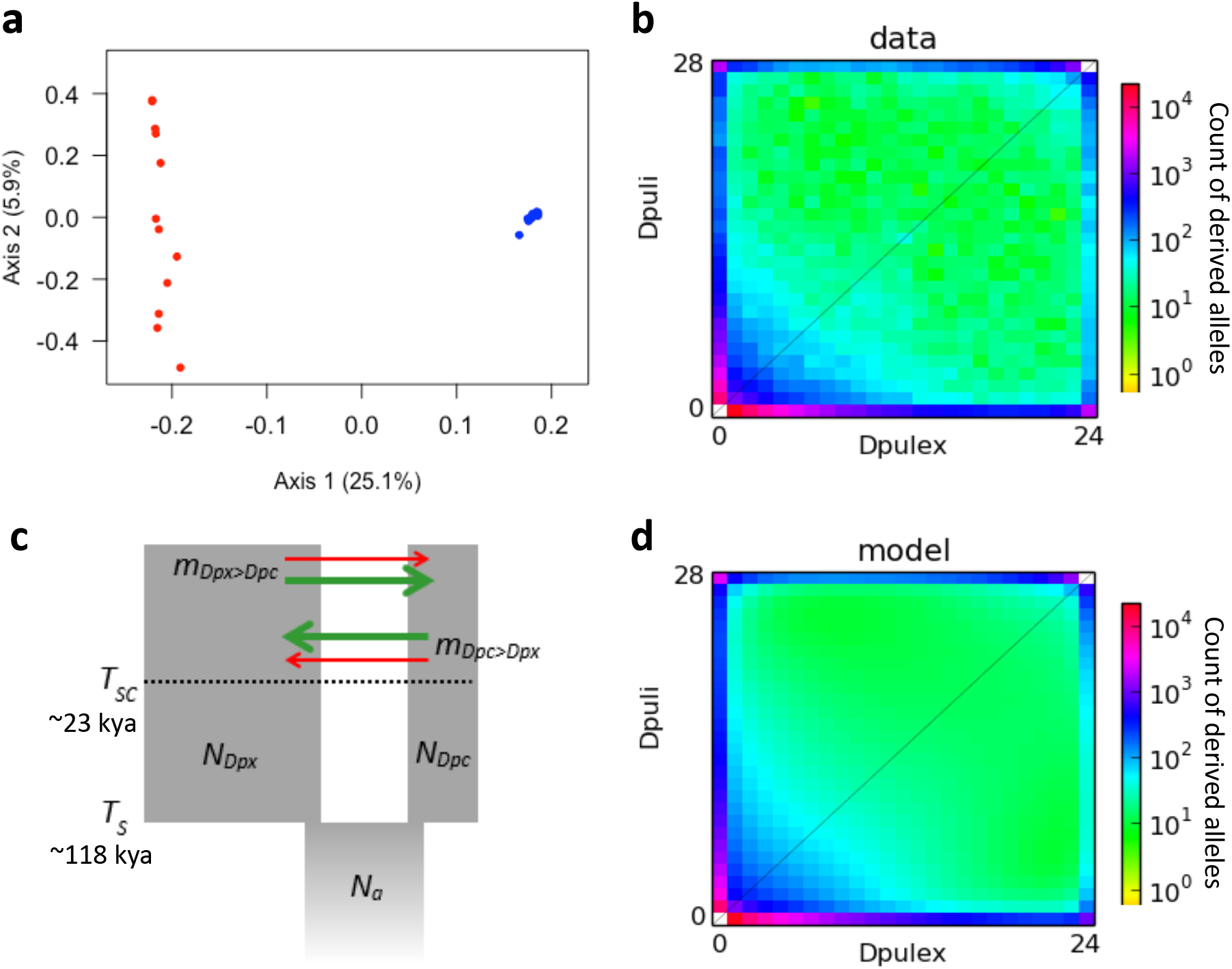
Demographic history of *D. pulicaria* and *D. pulex* speciation. **(a)** Principal component analysis of 118,274 neutral SNPs across 14 *D. pulicaria* (blue) and 12 *D. pulex* (red) natural isolates. **(b)** The unfolded joint allele-frequency spectrum (AFS) for *D.pulex* and *D.pulicaria* samples, showing the count of derived alleles for 118,274 four-fold degenerate neutral SNPs, polarized with *D. obtusa* as an ancestral outgroup. **(c)** The secondary contact with heterogeneous migration rate model (*SC2m*), including 10 parameters: the ancestral effective population size (*N_a_*), the *D. pulex* and *D. pulicaria* effective population sizes after splitting (*N_Dpx_* and *N_Dpc_*), the splitting time (*T_s_ circa* 118 kya) and the time since secondary contact (*T_SC_ circa* 23 kya). Migration rate from *D. pulex* into *D. pulicaria (m_Dpx>Dpc_*) and in the opposite direction (*m_Dpc>Dpx_*) can take two free values in proportions *p* (green arrows) and 1-*p* (red arrows). **(d)** The maximum-likelihood AFS obtained under the *SC2m* model.

This *SC2m* model and its maximum-likelihood parameters describe the evolutionary history of divergence in North American *D. pulex* and *D. pulicaria* (Table 1). Under this model, divergence accumulated during a period of isolation four times longer than the age of secondary contact. Using *de novo* mutation rate estimates from *Daphnia* mutation-accumulation experiments^33^ and estimates of the number of *Daphnia* generations per year^38,41^, we convert the divergence model parameters into historical time scales. While *D. pulex* reach peak fecundity in approximately half the time as *D. pulicaria*, they only reproduce in the spring/summer months while *D. pulicaria* can maintain reproduction year-round^41^. Accounting for these factors, we estimated both species produce an average of 7 generations per year for this analysis. *Daphnia pulex* and *D. pulicaria* are estimated to have diverged ~118,000 years ago (*T_s_* in Fig. 1c), with a ~95,000 year period of isolation followed by a ~23,000 year period of secondary contact with gene flow (*T_SC_* in Fig. 1c). Effective population size estimates revealed that *D. pulex* experienced significant population expansion (1.77 times higher than ancestral size) while *D. pulicaria* experienced population contraction (0.63 times lower than ancestral size). Furthermore, introgression following secondary contact was strongly asymmetric, with the rate from *D. pulex* to *D. pulicaria* four times as much as the reverse. The *SC2m* model indicated that a substantial proportion of the *D. pulicaria* and *D. pulex* genomes experienced highly reduced gene flow compared to the rest of their genomes, with rates over an order of magnitude less in introgression-resistant regions. Thus, the model describes heterogeneous divergence patterns between *D. pulex* and *D. pulicaria* that resulted from the erosion of divergence through differential introgression after secondary contact.

**Table 1.**
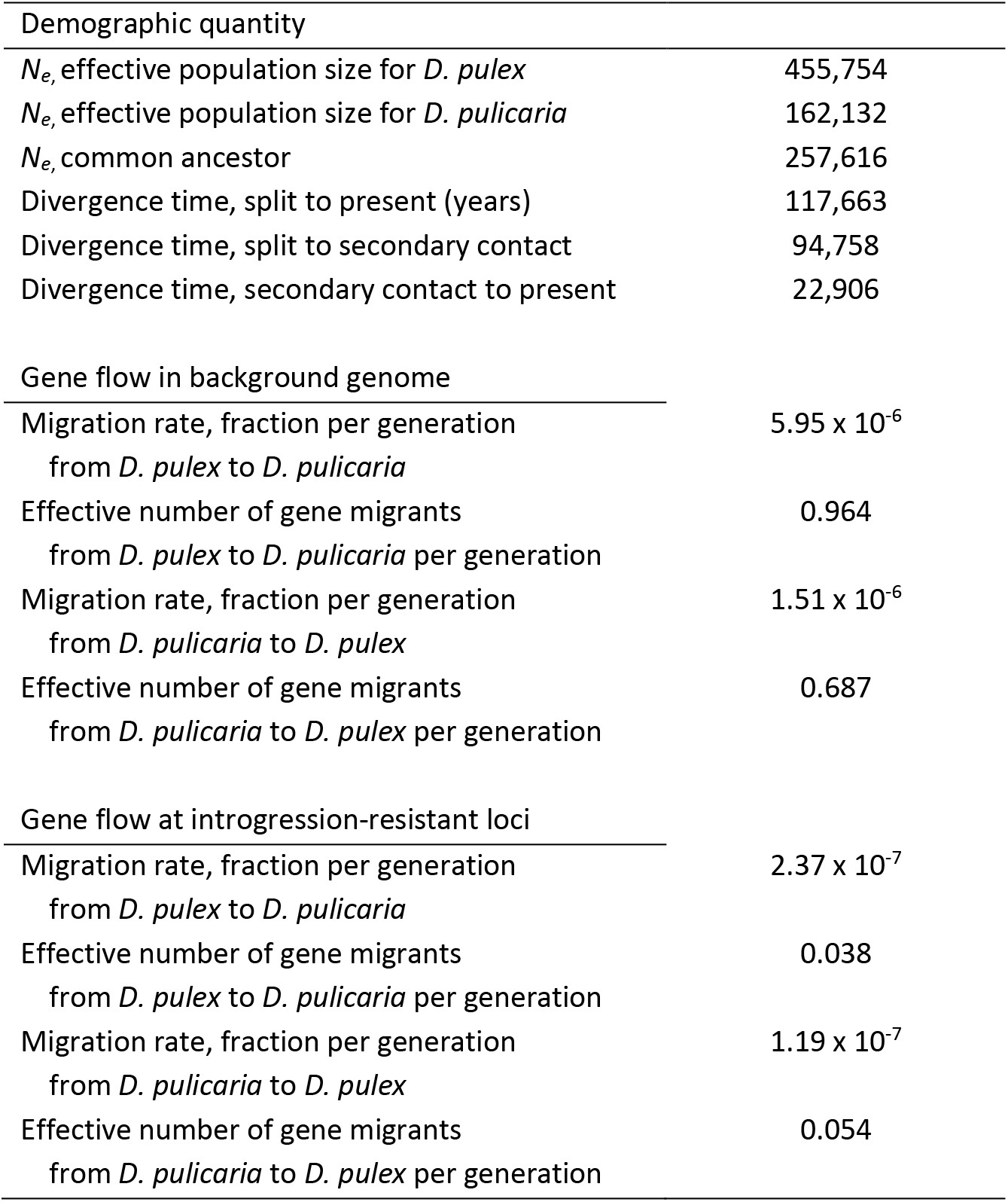
Maximum-likelihood demographic quantities from *SC2m* model for genome-wide divergence between *D. pulex* and *D. pulicaria*.

### Heterogeneous genomic landscape reveals large contiguous “continents of divergence”

We analyzed several measures of relative and absolute species divergence between *D. pulex* and *D. pulicaria* by applying genome scans in non-overlapping sliding windows across the genome. Utilizing the *D. pulicaria* chromosome-scale assembly, we were able to visualize the complete genomic landscape of divergence across all 12 *Daphnia* chromosomes (Fig. 2). Monomorphic fixed differences between species accumulate throughout the genome over time as mutations reach fixation in one species lineage due to genetic drift or selection. In total, we identified 73,652 fixed differences (3.59% of total SNPs) between the *D. pulex* and *D. pulicaria* samples. The distribution of fixed differences across the genome (measured as D_f_, the density of fixed differences) is remarkedly concentrated to certain regions of the genome (Fig. 2b). We identified 12 large contiguous regions (>1 Mb) containing 93.6% of all fixed differences, with most chromosomes containing a single large divergence region, while the background genome is mostly devoid of fixed differences between the species (Supplementary Table 4). These large genomic divergence regions encompass 36% of the total genome and range in size from 1.8 Mb to 6.7 Mb, with a median size of 5.3 Mb, and cover from ~16% to ~82% of their respective chromosomes (Supplementary Fig. 2). Due to the size, contiguity, and chromosome coverage of these divergence regions, we refer to these regions as genomic “continents of divergence” spread throughout the background genome. The genomic landscape of divergence supports the *SC2m* demographic model previously described, indicating a history of gene flow and introgression between the species, homogenizing the background genome, while the divergence continents are resistant to interspecific introgression and preserve accumulated fixed differences.

**Figure 2.**
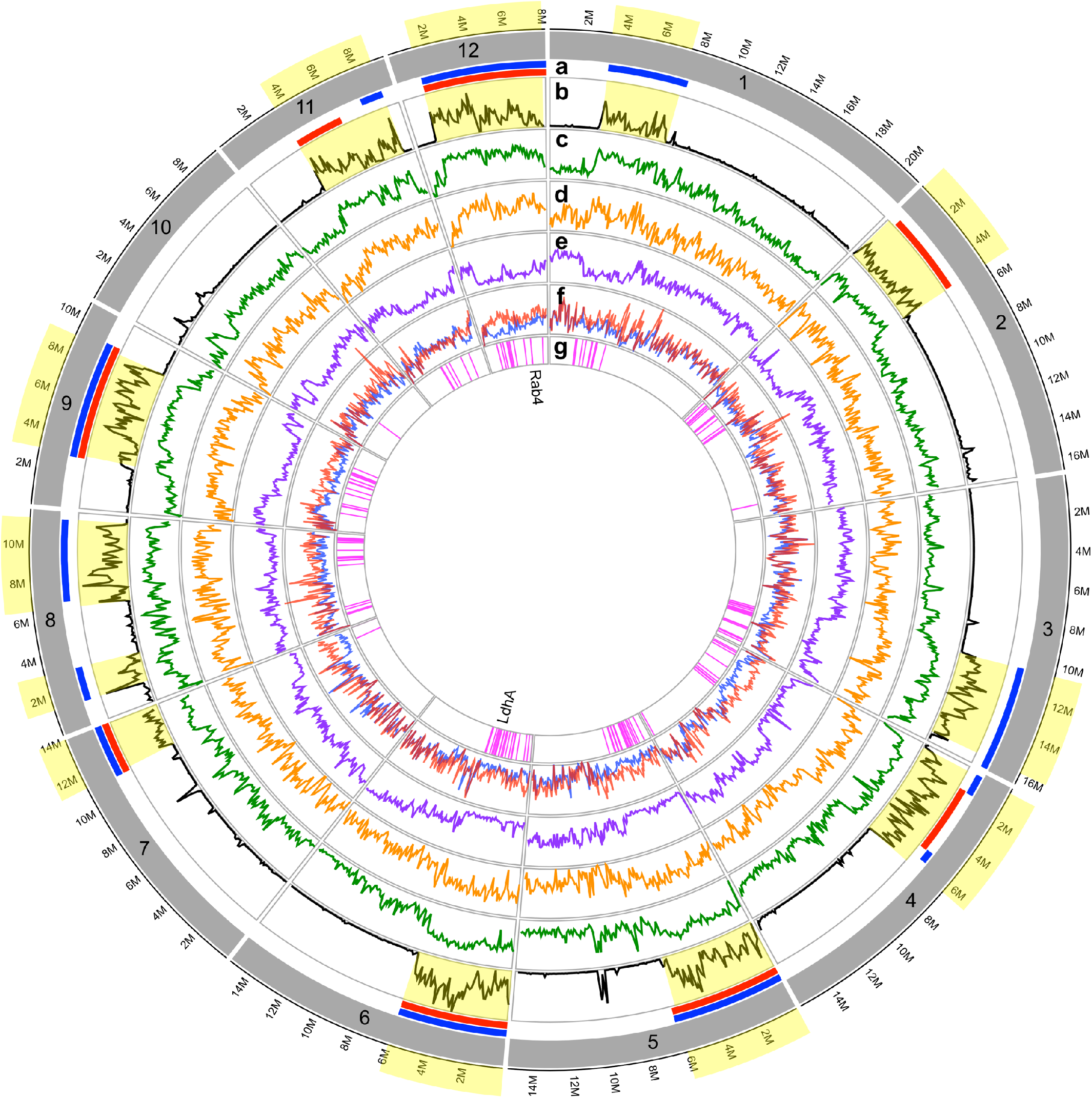
Heterogeneous genomic landscape of divergence in *Daphnia*. Large contiguous genomic regions spread across 11 *Daphnia* chromosomes show marked divergence between *D. pulicaria* and *D. pulex*. These “continents of divergence” account for 36% of the total *D. pulicaria* genome while containing 94% of the fixed differences between the species (highlighted in yellow). **(a)** Regions of chromosomal rearrangement occurring in the *D. pulicaria* lineage (blue), the *D. pulex* lineage (red), or both lineages. **(b-f)** 100 kb non-overlapping sliding windows across the genome comparing genetic statistics between *D. pulicaria* and *D.pulex* isolates: **(b)** Black - Density of fixed differences (D_f_), **(c)** Green – Relative divergence (F_ST_) **(d)** Orange - Absolute divergence (D_xy_), **(e)** Purple - Proportion of shared polymorphism (SNPs not unique to one species) **(f)** Nucleotide diversity (π) for *D. pulicaria* (blue) and *D.pulex* (red). **(g)** Genomic positions of 142 candidate adaptive genes with significant evidence of positive selection (McDonald-Kreitman test, p < 0.05). The LdhA and Rab4 loci conventionally used to genetically distinguish between *D. pulicaria* and *D. pulex* species are denoted within the chromosome 6 and chromosome 12 continents of divergence.

In the continents of divergence, mean relative divergence (F_ST_) is 2.08 times higher, mean absolute divergence (D_xy_) is 1.53 times higher, and the mean density of fixed difference (D_f_) is 23.2 times higher than in the background genome (Table 2, Fig 2b-e). Nucleotide diversity in *D. pulicaria* (π_*Dpulicaria*_) is 1.37 times higher in the background genome compared to the divergence continents, likely due to introgression from *D. pulex* in the background genome, as described in the *SC2m* demographic model (Table 2, Fig. 2f). Genome-wide distributions of relative (F_ST_) and absolute (D_xy_) divergence in 100 kb sliding windows also highlight the heterogeneous landscape of divergence between the divergence continents and background genome (Supplementary Fig. 3).

**Table 2.** Mean divergence statistics between *D. pulex* and *D. pulicaria* across the background genome and divergence continents.

|  | Background<br>Genome | Divergence<br>Continents |
| --- | --- | --- |
| Proportion of<br>Genome | 0.640 | 0.360 |
| $F_{ST}$ | 0.275 (0.004) | 0.571 (0.004) |
| $D_f (10^{-3})$ | 0.052 (0.006) | 1.206 (0.031) |
| $D_{xy} (10^{-3})$ | 3.940 (0.053) | 6.011 (0.062) |
| **Proportion of**<br>**Shared SNP**<br>**Variation** | 0.472 (0.004) | 0.206 (0.003) |
| $\pi_{\text{Dpulex}} (10^{-3})$ | 3.316 (0.045) | 3.228 (0.036) |
| $\pi_{\text{Dpulicaria}} (10^{-3})$ | 2.717 (0.038) | 1.977 (0.030) |

We further analyzed the allele frequencies of the 2,049,444 bi-allelic SNPs within and between *D. pulex* and *D. pulicaria* to document the amount of shared and species-specific polymorphism across the genomes. Overall, ~1.32 million SNPs (64.4%) are variable in one species only, with *D. pulex* having nearly twice the number of species-specific polymorphisms as *D. pulicaria*, ~874k (42.6%) vs ~446k (21.7%) SNPs respectively. Also, the amount of species-specific vs shared polymorphism is markedly different between the divergence continents and the background genome (Fig. 2e). In the divergence continents 20.2% of SNPs are shared, likely due to shared ancestral polymorphisms, while the remaining 79.8% of SNPs are unique to only one species. In contrast, the amount of shared polymorphism in the background genome is more than twice as high (46.6% shared), suggesting that interspecific introgression and incomplete lineage sorting affect the background genome more freely than the divergence continents.

### Continents of divergence contain numerous candidate adaptive genes

To determine the possibility of a subset of *Daphnia* genes being under positive selection, we evaluated the neutrality index (NI), defined to be the ratio of P_n_/P_s_ to D_n_/D_s_^48^. Values of NI < 1.0 indicate excess divergence of replacement substitutions and are generally taken to imply positive selection for adaptive amino acid substitutions. Comparing *D. pulicaria*-*D. pulex* gene divergence, we found NI < 1.0 for 1344 of the 2032 genes (66%) for which the computation could be carried out (which requires a nonzero denominator). Thus, there is evidence for excess amino acid sequence divergence in a fraction of *D. pulicaria/D. pulex* genes (~4.9% of total genes). We applied the McDonald-Kreitman test (M-K test) to the *D. pulicaria/D. pulex* genes, which measures the amount of adaptive evolution between species and estimates the proportion of substitutions that are adaptive (α)^49,50^. We identified 142 genes with significant evidence of positive selection (p < 0.05, mean α = 0.941 (0.005), mean NI = 0.088 (0.007)), which we denote as candidate adaptive genes of interest (Supplementary Table 5). These candidate adaptive genes are spread throughout the continents of divergence, ranging from 2 to 21 genes per chromosome, with an average of 13 candidate genes per chromosome (Fig. 2g).

In addition, we tested the top 1% most differentiated outlier F_ST_ windows (20 kb windows, F_ST_ > 0.788) for enrichment of functional pathways. The KEGG phototransduction – fly pathway was the only significantly enriched pathway (Fisher’s exact test, Adj. p-val = 0.033). Arthropods use rhabdomeric opsin proteins (r-opsins) for photoreception and vision, and this phototransduction signaling pathway has been annotated in *Drosophila*^51,52^. The candidate adaptive gene set includes four r-opsins with evidence of positive selection between *D. pulicaria* and *D. pulex*, and an additional four r-opsin genes were included in the F_ST_ outlier windows with phototransduction pathway enrichment.

### Large-scale chromosomal rearrangements are associated with continents of divergence

To investigate the role of chromosomal structure in speciation, we aligned the *D. pulicaria* and *D. pulex* reference genome assemblies and visualized chromosome synteny between the species. The whole-genome alignment revealed numerous structural rearrangements between the species (Fig. 3). Some chromosomal rearrangement regions are structurally complex and include multiple inversions and intrachromosomal translocations. These regions correspond with the continents of divergence identified in the population genomics analyses (Supplementary Table 4). Similar to the continents of divergence, rearrangements mostly form one large contiguous region per chromosome, covering ~40% of each chromosome on average, and ~35% of the total chromosome sequence. For example, the leading 5 Mb region of chromosome 2 contains a large intrachromosomal translocation where adjacent 1.4 Mb and 3 Mb segments have exchanged chromosomal order between the species, corresponding to the identified divergence continent (Fig. 3c). Moreover, chromosome 8 contains two continents of divergence separated by a 3.4 Mb homogenized region. We found multiple intrachromosomal translocations combined with inversions on chromosome 8 corresponding to these continents of divergence, while the intervening homogenized region is collinear (Fig. 3b). The chromosome 4 continent of divergence harbors a large inversion containing multiple nested inversions, indicating a history of chromosomal inversion events in the region (Fig. 3d). These findings suggest that chromosomal rearrangements have played an important role in the formation and preservation of interspecific divergence between *D. pulex* and *D. pulicaria*.

**Figure 3.**
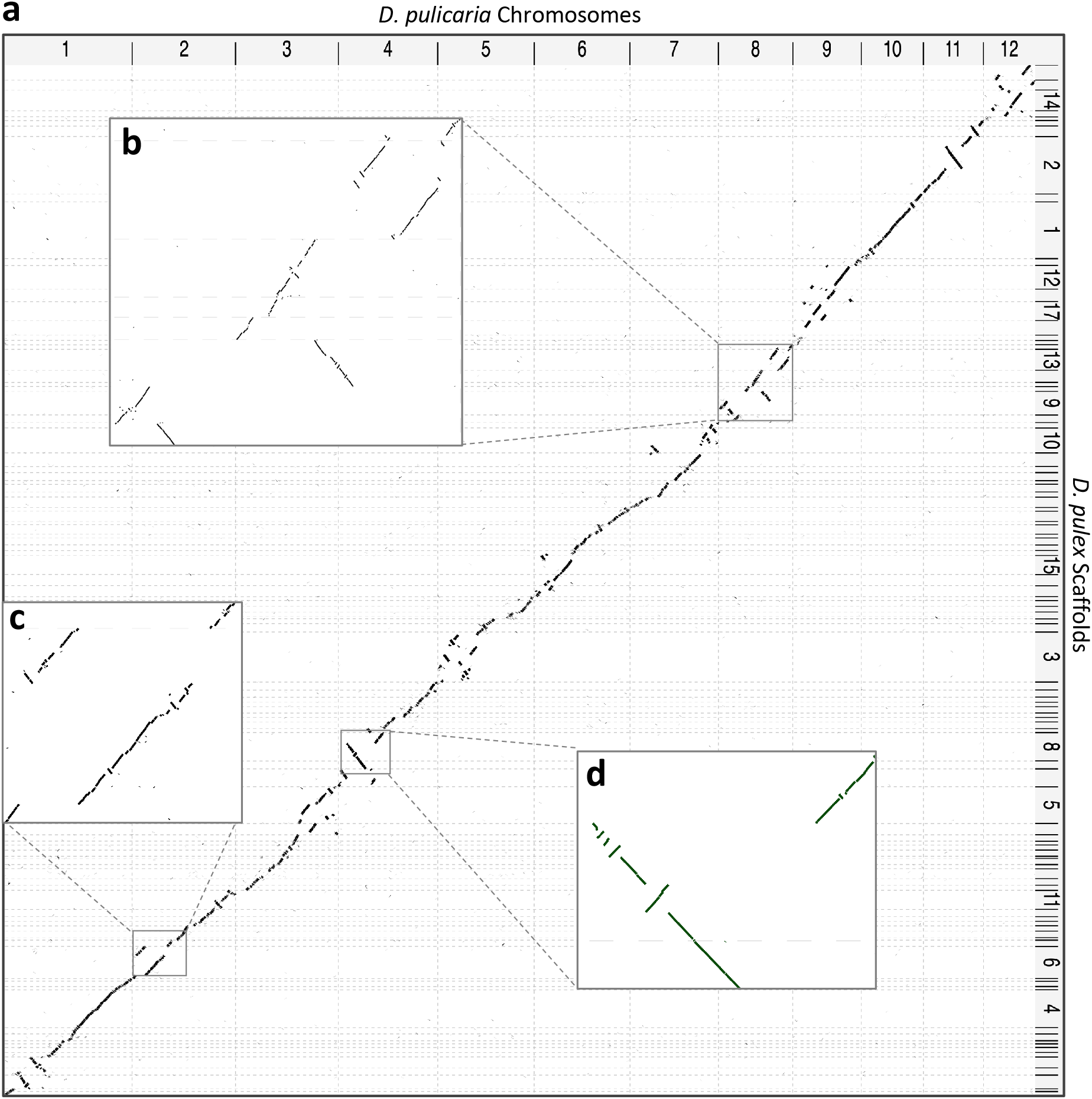
Genome synteny between *D. pulicaria* and *D. pulex* reveals numerous chromosomal rearrangements. **(a)** Dot plot visualization of *D.pulicaria-D.pulex* whole genome alignment between *D. pulicaria* chromosomes (x-axis) and *D. pulex* scaffolds (y-axis), revealing numerous inversions, translocations, and complex chromosome rearrangements. The main diagonal with a slope of 1 represents collinear regions of the genomes, while disruptions of this diagonal indicate regions of insertion, deletion, or translocation. Regions with an inverted slope of −1 represent inversions between the two genomes. Genomic regions containing chromosome rearrangements correspond with the large continents of divergence identified from population genomic analyses. **(b)** Chromosome 8 alignment showing intrachromosomal translocations and large inversions, corresponding with the two chromosome 8 divergence continents **(c)** Chromosome 2 alignment showing a large intra-chromosomal translocation corresponding with the continent of divergence spanning the first 5 Mb of chromosome 2. **(d)** Chromosome 4 alignment showing a large inversion with several smaller nested inversions contained within the inversion, corresponding with the continent of divergence spanning the first 6 Mb of chromosome 4.

To better understand the evolutionary trajectory of these chromosomal rearrangements, we used the genome assembly of the outgroup species *D. obtusa* to infer the ancestral genome structure and determine in which *Daphnia* lineage chromosomal rearrangements occurred. Genome synteny between *D. pulicaria – D. pulex, D. pulicaria – D. obtusa*, and *D. pulex – D. obtusa* was analyzed to determine collinear or rearranged chromosomal regions in *D. pulicaria* and *D. pulex*, relative to *D. obtusa* (Supplemental Fig. 4). Collinear sequences were inferred to be the ancestral genome structure, while rearranged genome structure indicates the lineage(s) containing chromosomal rearrangements. We determined that three chromosomal rearrangement regions were unique to the *D. pulicaria* lineage (on chromosomes 1, 3 and 8), one region was unique to the *D. pulex* lineage (on chromosome 2), while the other seven regions showed a history of chromosomal rearrangement in both species (Fig. 1a, Supplemental Table 4). Chromosome 1 synteny indicates an ~4.4 Mb inversion in the *D. pulicaria* lineage, while the *D. pulex* and *D. obtusa* lineages are collinear, indicating the ancestral genome structure. The chromosome 2 intrachromosomal translocation (see Fig. 3c) is unique to the *D. pulex* lineage, while *D. pulicaria* and *D. obtusa* lineages are collinear. While chromosome 1 and chromosome 2 contained rearrangements unique to a single lineage, multi-species synteny indicates the majority of chromosomal rearrangement regions are the result of multiple chromosomal events occurring in both *D. pulicaria* and *D. pulex* lineages. For instance, chromosome 11 contains two large inversions between *D. pulicaria* and *D. pulex*, and multi-species synteny indicates that one of the inversions occurred within the *D. pulicaria* lineage while the other occurred within the *D. pulex* lineage, together forming a continent of divergence spanning over half of chromosome 11 (Fig. 4). Other chromosomal rearrangement regions, such as on chromosomes 5, 6, 7, 9, and 12, show complex syntenic order between species, indicating an evolutionary history of multiple inversions or complex rearrangements occurring in both *D. pulicaria* and *D. pulex* lineages.

**Figure 4.**
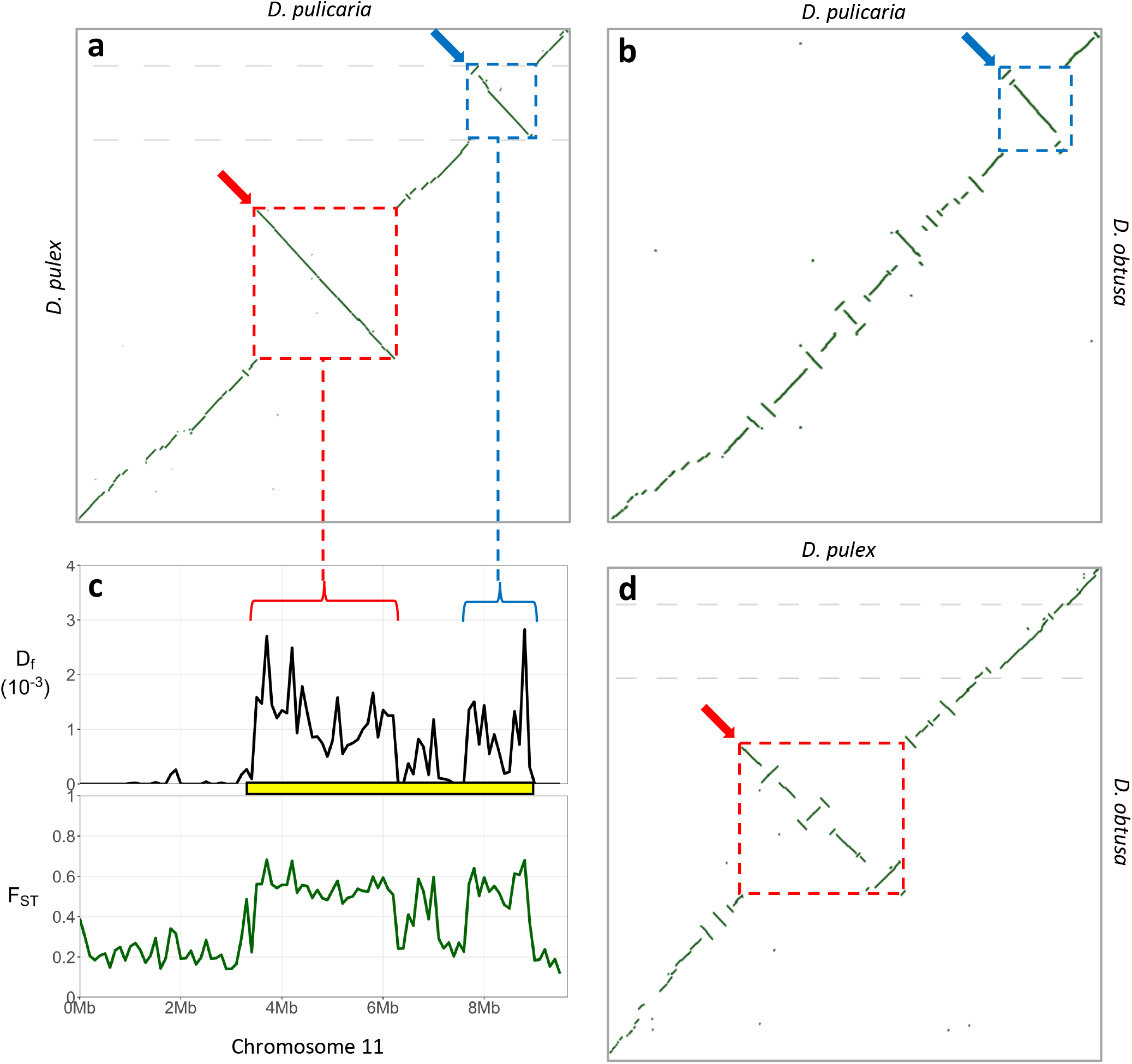
Chromosomal inversions in both *D. pulicaria* and *D. pulex* lineages are associated with divergence. Synteny between three *Daphnia* species across chromosome 11 reveals two large inversions between *D. pulicaria* and *D. pulex* associated with *D. pulicaria-D.pulex* divergence, while numerous smaller inversions exist amongst multiple *Daphnia* lineages. Synteny with the outgroup species *D. obtusa* resolves the ancestral genome orientation and establishes the *Daphnia* lineage for the two large inversions. A 2.7 Mb inversion in the *D. pulex* lineage (red dashed lines/arrow) is associated with the first half of the chromosome 11 divergence continent (yellow block) while a 1.2 Mb inversion in the *D. pulicaria* lineage (blue dashed lines/arrow) is associated with the end of the divergence continent. **(a)** *D. pulicaria* – *D. pulex* synteny dotplot across chromosome 11 shows two large inversions. **(b)** *D. pulicaria – D. obtusa* synteny dotplot across chromosome 11 indicates the second inversion (blue dashed lines/arrow) established in *D. pulicaria* lineage. **(c)** Divergence measures density of fixed differences (D_f_, black) and fixation index (F_ST_, green) across chromosome 11 show continent of divergence (yellow block) between *D. pulicaria* and *D. pulex*. **(d)** *D. pulex* – *D. obtusa* synteny dotplot across chromosome 11 indicates the first inversion (red dashed lines/arrow) established in *D. pulex* lineage.

## Discussion

Genomic sequence variation, allelic and structural alike, provides the substrate for evolution. Here we leverage the new chromosome-scale assembly of *D. pulicaria* and improved *D. pulex* assembly, as well as natural population resequencing, to characterize the role that chromosomal rearrangements have contributed to genomic divergence in the context of ecological speciation. We describe the landscape of divergence at the genome-wide scale, reconstruct the evolutionary history of speciation, and uncover the important role of chromosomal rearrangements in preserving adaptive divergence.

The suggestion that structural rearrangements can be important to adaptation and speciation is not new^1,6^. In this study we highlight genome-scale empirical evidence for the role of chromosomal rearrangements in ecological speciation with gene flow. Rieseberg argued that chromosome rearrangements reduce gene flow more by suppressing recombination than by reducing fitness during speciation and suggested that these rearrangements facilitate speciation in the presence of gene flow, a process called the suppressed recombination model of speciation^6^. Kirkpatrick and Barton extended this model, showing through simulations that chromosome inversions capturing multiple locally adapted alleles can spread to high frequency and suppress recombination between hybridizing populations or species^9^. Our results strongly support this suppressed recombination model. We identify chromosomal rearrangements harboring multiple locally favored habitat-associated alleles and divergence patterns suggesting that these rearrangements limited introgression during secondary contact, likely due to chromosomal incompatibilities and deleterious effects. Divergence among *Timema* ecotypes suggested that large inversions are more likely to underlie adaptive divergence than other structural variants^13^. While we find that large inversions are associated with adaptive divergence, we also find that intrachromosomal translocations are prevalent, with both often forming complex rearrangement regions associated with adaptive divergence between *D. pulex - D. pulicaria*.

Chromosomal speciation with gene flow between *D. pulex – D. pulicaria* generated a heterogeneous genomic landscape consisting of continents of divergence spread throughout the genome. The field of population genomics has previously focused attention on speciation in which divergent selection acts on only a handful of genes that reside in just a few small isolated chromosomal regions, giving rise to the metaphor of ‘genomic islands of speciation’^15,23,53^. Since then, a wide range of speciation dynamics have been explored, including many differentiation patterns and underlying speciation processes, increasingly incorporating whole genome sequencing^19^. Our *Daphnia* speciation results contribute to this growing field and provide new empirical evidence of widespread continents of genomic divergence maintained by chromosomal rearrangements. While other researchers have identified chromosomal rearrangements correlated with divergence, such as in *Gasterosteus* stickleback ecotypes^11^ and *Heliconius* butterfly morphs^54^, they only represented a small proportion of the genome. We show that chromosomal rearrangements associated with divergence in *Daphnia* encompass more than a third of the entire genome across nearly all chromosomes.

The continents of divergence harbor genes that provide adaptive advantages for the distinct habitat of each species. We identify 142 genes showing significant signatures of positive selection in *D. pulex – D. pulicaria* evolution. Functional analyses show several of these genes correspond to observed phenotypic and habitat differences, including photoperiod response and phototaxis, growth regulation, metabolism, and longevity. For instance, we identified evidence of selection in a *Daphnia* homolog of the *Drosophila melanogaster* gene I’m not dead yet (*Indy*). Mutations in the *Indy* gene and homologs have been shown to affect metabolic state and longevity in *D. melanogaster*, *Caenorhabditis elegans*, and *Mus musculus*^55^. Interestingly, *D. pulicaria* can have more than double the longevity of *D. pulex* in proper environment conditions^41^.

Several genes known to regulate and contribute to the phototransduction pathway show evidence of positive selection and differentiation between *D. pulex-D. pulicaria. Daphnia pulex* has the largest known family of opsins due to extensive duplication of opsin genes in the *Daphnia* lineage, including 29 r-opsins^29,56,57^. Phylogenetic and functional analyses of *Daphnia* r-opsins suggest that diverse r-opsins are maintained due to diverse functional roles in photoreception and vision^56^. In total, we identified 8 highly differentiated r-opsins between *D. pulex* and *D. pulicaria*. In addition, the transcription factor Kruppel (*Kr*) and the retinal degeneration B (*rbgB*) gene both had evidence of positive selection. Kruppel is enriched in the *Drosophila* eye and controls photoreceptor differentiation and regulates r-opsin expression^58^, while retinal degeneration B (*rbgB*) gene encodes an integral membrane protein that is essential for phototransduction and prevention of retinal degeneration in *Drosophila^59,60^*. Light attenuation is very different between pond and lake habitats, and therefore *D. pulex* and *D. pulicaria* have different photoperiodic responses^35^, which may explain why the phototransduction pathway would be under such strong selection. Our results suggest that the two species have evolved these different responses to light through their r-opsin gene repertoire and various membrane, signaling, and regulatory genes within the phototransduction pathway.

The *Ldh*A gene has historically been used as a genetic marker to distinguish *D. pulex* and *D. pulicaria* due to fixed substitutions affecting the electrophoretic mobility of species-specific alleles^37,61^. For this reason, researchers have suggested the possibility that *Ldh* variation is adaptive, but this hypothesis had not yet been tested^62^. The candidate adaptive gene Kruppel is located just 62 kb downstream from the *Ldh*A gene within the Chromosome 6 divergence continent. While we also identified the previously reported species-specific alleles of *LdhA*, we did not find evidence that *Ldh* variation is adaptive. Instead, it appears the *Ldh* divergence is due to tight linkage with candidate adaptive genes in the Chromosome 6 continent of divergence. To a lesser degree, the *Rab4* gene has also been used as a genetic marker for *D. pulex* and *D. pulicaria*^30^. *Rab*4 is located within the Chromosome 12 divergence continent with 14 candidate adaptive genes, including a r-opsin gene just 43 kb upstream of *Rab4*.

Demographic modeling of speciation can link the evolutionary genome landscape with ecological or geological changes driven by historical environmental changes. The maximum-likelihood demographic model for the ecological partitioning of these two species is consistent with geological changes driven by historical climatic changes. We provide evidence that the two species diverged under a mixed geographic mode of evolution. Our ability to capture the effects of variable gene flow that generate heterogeneous patterns of divergence was important to more accurately model *Daphnia* chromosomal speciation. Most likely, the initial divergence of the two species occurred around the end of the Eemian interglacial (~115 kya) and beginning of the last glacial period ^63^, after which the two species subsequently adapted to different ecological habitats in isolation over a ~95,000 year period. The two species were previously estimated to have diverged ~82 kya (95% HPD interval: 58 kya – 126 kya) using an IM model, but only six loci were sequenced and mutation rates from *Drosophila* used^64^. In this study we use over 100,000 genome-wide neutral markers and our divergence age estimates are based on measured mutation rates from *D. pulex* mutation accumulation experiments^33^. Most of the sampled aquatic habitats where we collected the *Daphnia* isolates for this study were completely glaciated during the Last Glacial Maximum^65^. Secondary contact and subsequent gene flow is estimated to occur after the last glacial maximum in North America^66^, as glaciers retreated and the species colonized the newly created aquatic habitats. While computer simulations have previously shown that the mixed geographic mode of evolution can generate divergent chromosomal rearrangements that capture locally favored alleles^10^, we now provide strong genomic evidence of this evolutionary phenomenon occurring during *Daphnia* ecological speciation.

*Daphnia* has a long history as model for ecological and evolutionary research^67–69^. While earlier genome assemblies have advanced our understanding of environmental genomics^29,45^, their fragmentation has limited studies of genome structural variation and architecture. The chromosome-scale assemblies generated for this study allowed us to establish a role of chromosomal rearrangements in adaptive divergence in *Daphnia*, providing new genome-scale empirical support for chromosomal speciation. These results show that large-scale chromosomal rearrangements helped preserve adaptive divergence within the two *Daphnia* species experiencing gene flow during secondary contact, generating a heterogeneous landscape of divergence.

## Methods

### Genome Assembly

For the *D. pulicaria* assembly, a natural isolate was collected from Lake Sixteen, Martin Township, Michigan (LK16) and inbred one generation in the lab to reduce heterozygosity for improved assembly. A natural isolate from Portland Arch, Indiana (PA42) was used for the *D. pulex* assembly. The *D. pulicaria, D. pulex* and *D. obtusa* reference genomes were assembled using a combination of 80x-90x PacBio SMRT long-read sequencing, Illumina long-insert mate pair sequencing (2kb, 5kb, 10kb), and chromosome conformation capture sequencing (Dovetail Chicago and Hi-C). First, the PacBio reads were assembled using the Pacific Biosciences hierarchical genome-assembly process (HGAP ver. 4)^70^ with the Falcon assembler^71^. The PacBio contigs were then scaffolded with the Illumina mate pair sequences using the BOSS scaffolder^72^. Dovetail Chicago in vitro proximity ligation sequencing was generated for both *D. pulicaria* and *D. pulex*, and was used with the HiRise proximity ligation genome scaffolding software^73^ to further improve the assembly contiguity. The *D. pulicaria* and *D. obtusa* genomes were further scaffolded into chromosome-scale assemblies using Dovetail Hi-C long-range proximity ligation. The final assembled chromosome-scale scaffolds were numbered from largest to smallest (1 being the largest and 12 the smallest assembled chromosome). Microsatellite markers from the first *D. pulex* genetic linkage map^74^ were mapped the *D. pulicaria* chromosomes in order to match the *D. pulicaria* chromosomes with the previously described *D. pulex* linkage groups (Supplemental Table 6).

### Genome Annotation

Genome assemblies were annotated with Funannotate pipeline (version 1.5.0, https://github.com/nextgenusfs/funannotate) using transcript evidence from Trinity *de novo* RNA-seq assemblies and HISAT2 mapping (Grabherr et al. 2011; Kim et al. 2019). We exposed *Daphnia* to 18 °C, 26 °C, NaCl, pH, Ni, UV, and Atrazine conditions, isolated RNA from the exposures, and sequenced using Illumina RNA-seq. In addition, we used previously published RNA-seq datasets from *Daphnia* developmental, defense response, and environmental sex determination studies^77,78^. The Funannotate pipeline ran as follows: (i) repeats were identified using RepeatModeler and soft masked using RepeatMasker, (ii) protein evidence from a UniProtKB/Swiss-Prot-curated database and 8 arthropod protein sets was aligned to the genomes using diamond v0.8.38^79^ and exonerate v2.4, (iii) transcript evidence was aligned using minimap2^80^ and PASA, (iv) *ab initio* gene predictors AUGUSTUS v3.3^81^ and GeneMark-ES v3.52^82^, (vi) consensus protein coding gene models were predicted using EvidenceModeler^83^. Functional annotation used available databases and tools, including PFAM, InterProScan, UniProtKB, KEGG ortholog pathways, eggnog-mapper, and PANNZER^84^.

### Population Genomic Analyses

From the NCBI Short Read Archive we downloaded previously published whole-genome sequences of 12 *D. pulex* isolates and 14 *D. pulicaria* isolates that are distributed across their known distribution range in North America. These isolates have been sequenced with 100bp or 150bp paired-end reads on Illumina platforms in previous studies^85–87^. Raw reads were trimmed and quality filitered using Trimmomatic v0.36 (parameters: ILLUMINACLIP:TruSeq3-PE-2.fa:2:30:10 LEADING:20 TRAILING:20 MINLEN:50). Trimmed reads were mapped to the *D. pulicaria* genome assembly using BWA-MEM v0.7.10 with default mapping parameters^88^ and variants were called with freebayes v1.2.0^89^. Bi-allelic SNPs with site quality > 40 and minor allele frequency > 0.05 were used for population genomic analyses, resulting in 2,049,444 final SNPs. Maximum likelihood phylogenies were created using RAxML v8^90^ and principal component analyses was done using SNPRelate R package^91^. Population genetic statistics across the genome (i.e. sliding windows) were computed using evo (Malinsky et al. 2018; https://github.com/millanek/evo) and vcftools^93^.

### Demographic Inference

The demographic history of *D. pulicaria* and *D. pulex* were inferred using a custom version of the software *dadi* v1.7^47^. We first identified 118,274 four-fold degenerate synonymous SNPs from the population genomics dataset, which we consider as neutral mutations for the demographic analyses. The genome sequence of a closely related species *D. obtusa* was used as an outgroup to the polarize the ancestral state of the neutral mutations. Thus, we produced the unfolded joint site-frequency spectrum (JSFS) for 14 *D. pulicaria* and 12 *D. pulex* isolates, a bidimensional matrix giving the number of counts of derived mutations segregating at x copies in *D. pulex* and y copies in *D. pulicaria*. We considered the four basic models representing alternative modes of divergence: Strict Isolation (SI), Isolation with Migration (IM), Ancient Migration (AM) and Secondary Contact (SC). In these models, an ancestral population of size N_ref_ splits into two populations of size N_1_ (*D. pulicaria*) and N_2_ (*D. pulex*) during T_S_ (SI and IM), T_AM_ + T_S_ (AM) or T_S_ + T_SC_ (SC) generations. Migration events during periods of T_S_ (IM), T_AM_ (AM) and T_SC_ (SC) generations occurred at rates *m*_12_ from *D. pulex* to *D. pulicaria*, and vice versa ^94^. In addition, we incorporated extensions to the models with gene flow in order to capture the effect of reduced gene flow around loci associated with adaptive divergence (heterogeneous gene flow: *2m*). These semi-permeability models allowed quantifying the proportion *P* of loci freely exchanged at rate *m*, whereas a proportion 1-*P* corresponded to the fraction of the genome experiencing a reduced effective migration rate *me*_12_ from *D. pulex* to *D. pulicaria*, and vice versa^26^. We also included Tine et al.’s optimization procedure improvements for *dadi*, including two rounds of simulated annealing followed by BFGS optimization to better identify the optimum model parameters. Python scripts to run these demographic models and optimization procedures are provided by Rougeux et. al. and available on GitHub (https://github.com/crougeux/Dadi)^27^. We generated 24 independent optimization runs for each model in order to check for model convergence and kept the best-fit run of each model in order to perform model comparisons on the basis of the Akaike information criterion (AIC).

### Candidate Speciation Genes

To detect signatures of positive selection in the genomes of *D. pulex* and *D. pulicaria*, we calculated the neutrality index (NI), defined to be the ratio of P_n_/P_s_ to D_n_/D_s_, and performed the McDonald-Kreitman test (MKT) for all eligible genes^48,49^. The snpEff software^95^ was used to annotate the functional context of all SNPs. Polymorphisms (P) and fixed divergences (D) for synonymous substitutions (designated as P_s_ and D_s_, respectively) and nonsynonymous substitutions (P_n_ and D_n_) were calculated. Fisher’s exact test was conducted on the 2×2 contingency table consisting of the synonymous/nonsynonymous polymorphisms and divergences for each eligible gene. Moreover, we estimated the proportion of divergences driven to fixation by positive selection (α) using the formula 1 – D_s_P_n_ / D_n_P_s_^50^.

### Whole Genome Alignment

The largest 150 *D. pulex* scaffolds (88.9% of total *D. pulex* assembly) were mapped onto the 12 *D. pulicaria* chromosome scaffolds to generate whole genome alignments using several alignment packages: MashMap^96^, Minimap2^80^, MUMmer^97^, and Mauve^98^. All four alignment packages revealed chromosomal rearrangements between the *D. pulicaria* and *D. pulex* genomes in similar chromosomal regions. We visualized the MashMap whole genome alignment using the D-GENIES dot plot software^99^ and manually curated the chromosomal rearrangement regions on the *D. pulicaria* chromosome assembly using alignment coordinates and genome visualization. MashMap was additionally used to map the *D. pulicaria* and *D. pulex* assemblies to the *D. obtusa* assembly to infer the ancestral genome structure and determine lineages for chromosomal rearrangements.

## Data Availability

The *D. pulicaria* (LK16 isolate), *D. pulex* (PA42 isolate), and *D. obtusa* (FS6 isolate) genome assemblies used in this study can be accessed at NCBI under the BioProject accessions PRJNA686130, PRJEB46221, PRJNA598691, and GenBank assembly accessions GCA_017493165.1, GCA_911175335.1, GCA_016170125.1. The raw reads used for the population genomic analyses can be accessed at NCBI Short Read Archive under the BioProject accessions PRJNA292080 and PRJNA351263, and sample accession numbers SAMN02252729–SAMN02252752, and SAMN06005639. Additional raw reads used for the outgroup species *D. obtusa* can be accessed at NCBI Short Read Archive under the BioProject accession PRJEB17737.

## Acknowledgements

Funding for this project was provided by the National Institutes of Environmental Health Sciences to J. Shaw (R01ES019324), the Office of the Vice President for Research at Indiana University, the National Science Foundation to M. Lynch (IOS-1922914), and the National Institutes of Health to M. Lynch (R35-GM122566-01). We thank S. Glaholt and S. Reynolds for *Daphnia* culturing, C. Caceres for *Daphnia* collection, and North Carolina State University Genomic Sciences Laboratory, Dovetail Genomics, and BGI Genomics for DNA sequencing. The authors acknowledge the Indiana University Pervasive Technology Institute and National Center for Genome Analysis Support for providing supercomputing and storage resources that have contributed to the research results reported within this paper. This research is based upon work supported by the National Science Foundation under Grant Nos. DBI-1062432 2011, ABI-1458641 2015, ABI-1759906 2018, and CNS-0521433 to Indiana University. Any opinions, findings, and conclusions or recommendations expressed in this material are those of the authors and do not necessarily reflect the views of the National Science Foundation, the National Center for Genome Analysis Support, or Indiana University. This research was supported in part by the Indiana Genomics Initiative and Lilly Endowment, Inc., through its support for the Indiana University Pervasive Technology Institute.

## Contributions

C.J., S.X, J.S and M.P. designed this project; C.J and Z.Y. assembled the genomes; C.J. analyzed population sequencing data and genome synteny; C.J., S.X., Z.Y., M.P., M.L., J.C., and J.S. interpreted data and contributed to writing the manuscript.

