## Supplemental Figures for "Chromosomal rearrangements preserve adaptive divergence in ecological speciation"

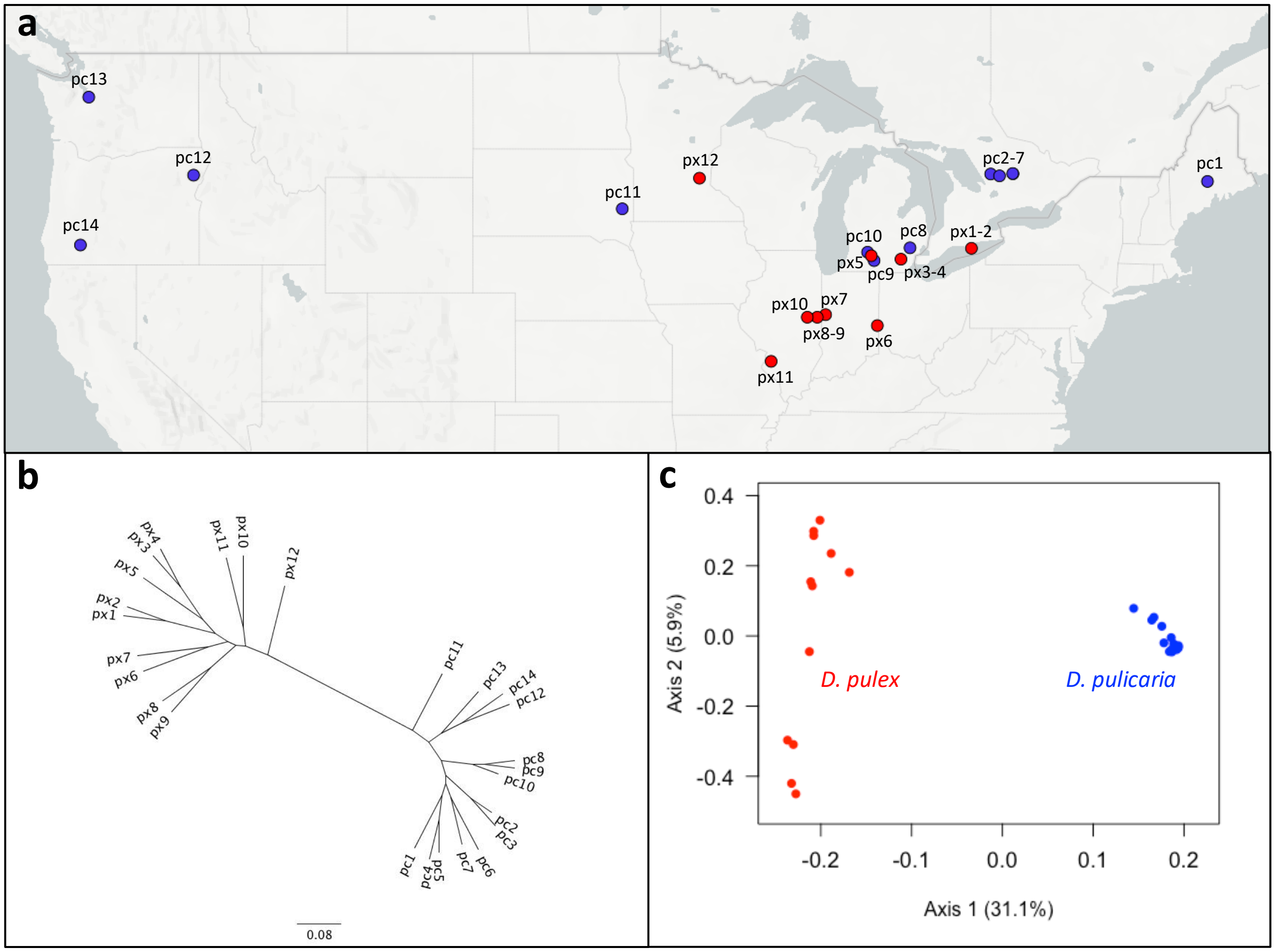


**Supplementary Figure 1. Geographic sampling and population structure of North American *D. pulex* and *D. pulicaria***
**(a)** Geographic location of 14 *D. pulicaria* (*pc*, blue) and 12 *D. pulex* (*px*, red) sampled across North America which were whole-genome sequenced and used for population genomic analyses. *D. pulicaria* were collected from lakes, while *D. pulex* were collected from vernal ponds. Samples are numbered from east to west (see Supp. Table 1). **(b)** Maximum likelihood unrooted phylogenetic tree of *D.pulicaria* and *D.pulex* samples based on ~2.05 million nuclear SNPs, showing *D. pulicaria* and *D. pulex* forming two distinct clades. **(c)** Principal component analysis of 2.05 million *D. pulicaria* and *D. pulex* SNPs. The first principal component, which separates *D.pulicaria* and *D.pulex* samples, explains 31.1% of the total variation in the dataset.

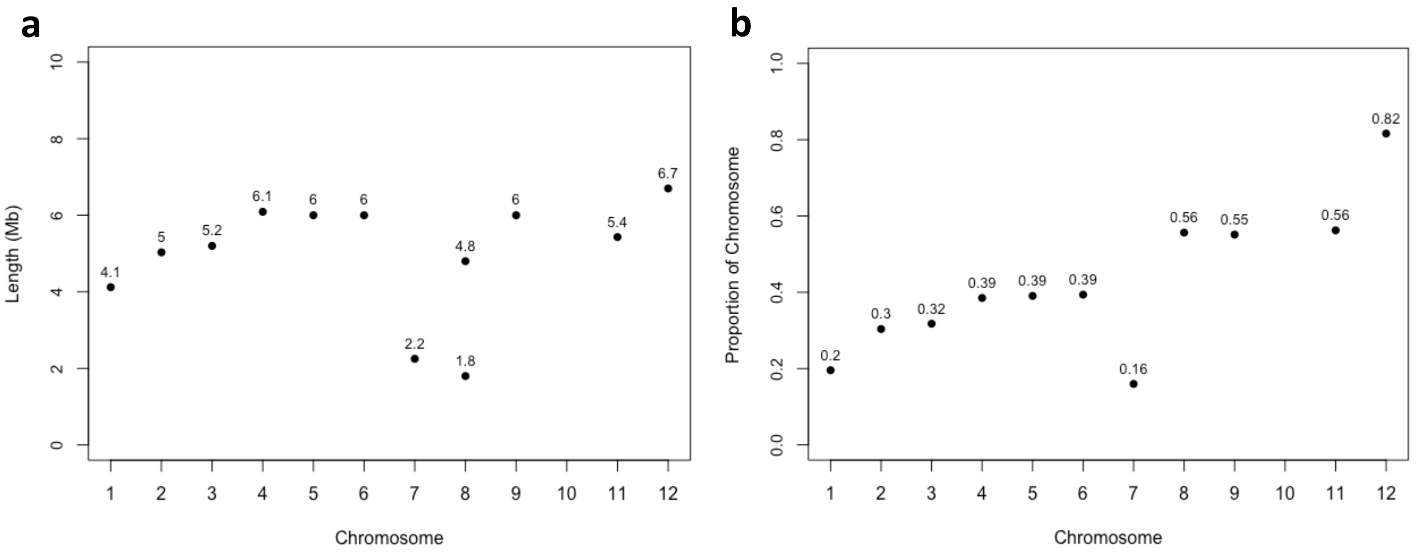


**Supplementary Figure 2. Length and chromosome span of 12 large contiguous genomic continents of divergence**
**(a)** Sequence **l**ength (Mb) of the genomic continents of divergence on each chromosome. **(b)** Proportion of the chromosome that each continent of divergences spans on its respective chromosome.

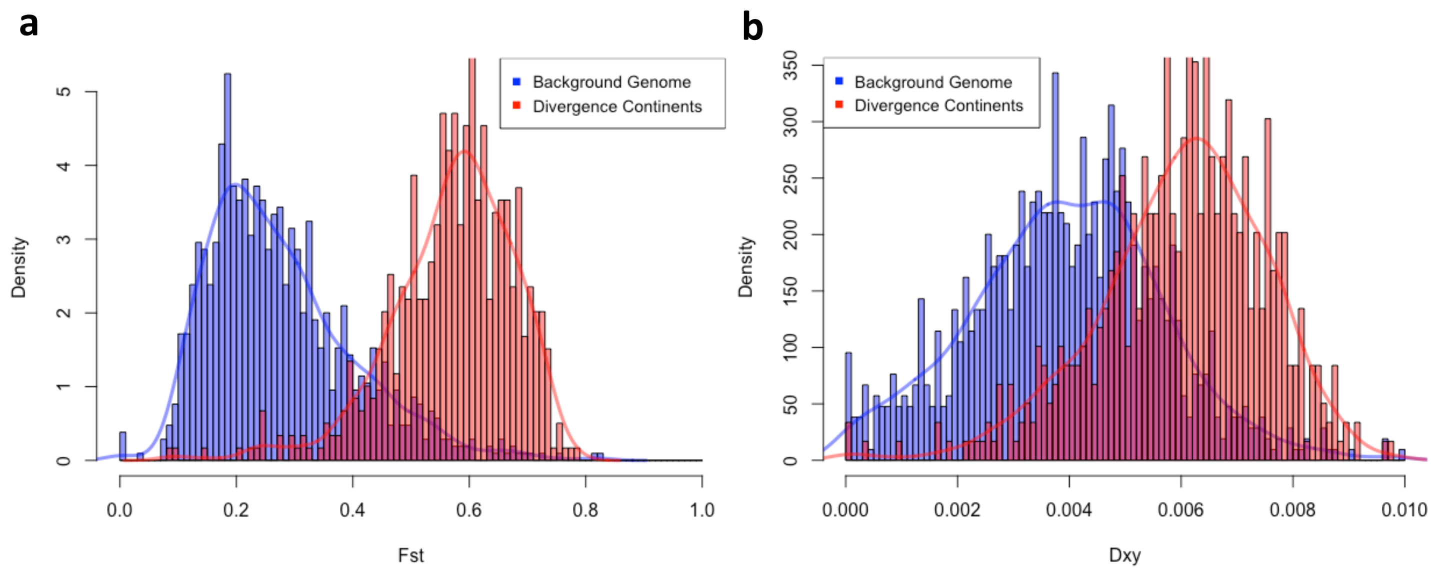


**Supplementary Figure 3. Distribution of relative ($F_{ST}$) and absolute ($D_{xy}$) divergence between *D. pulex* and *D. pulicaria***
**(a)** Distribution of relative divergence ($F_{ST}$) between *D. pulex* and *D. pulicaria* across 100 kb sliding windows in the background genome (blue) and the continents of divergence (red), with overlaid kernel density estimations. **(b)** Distribution of absolute divergence ($D_{xy}$) between *D. pulex* and *D. pulicaria* across 100 kb sliding windows in the background genome (blue) and the continents of divergence (red), with overlaid kernel density estimations.

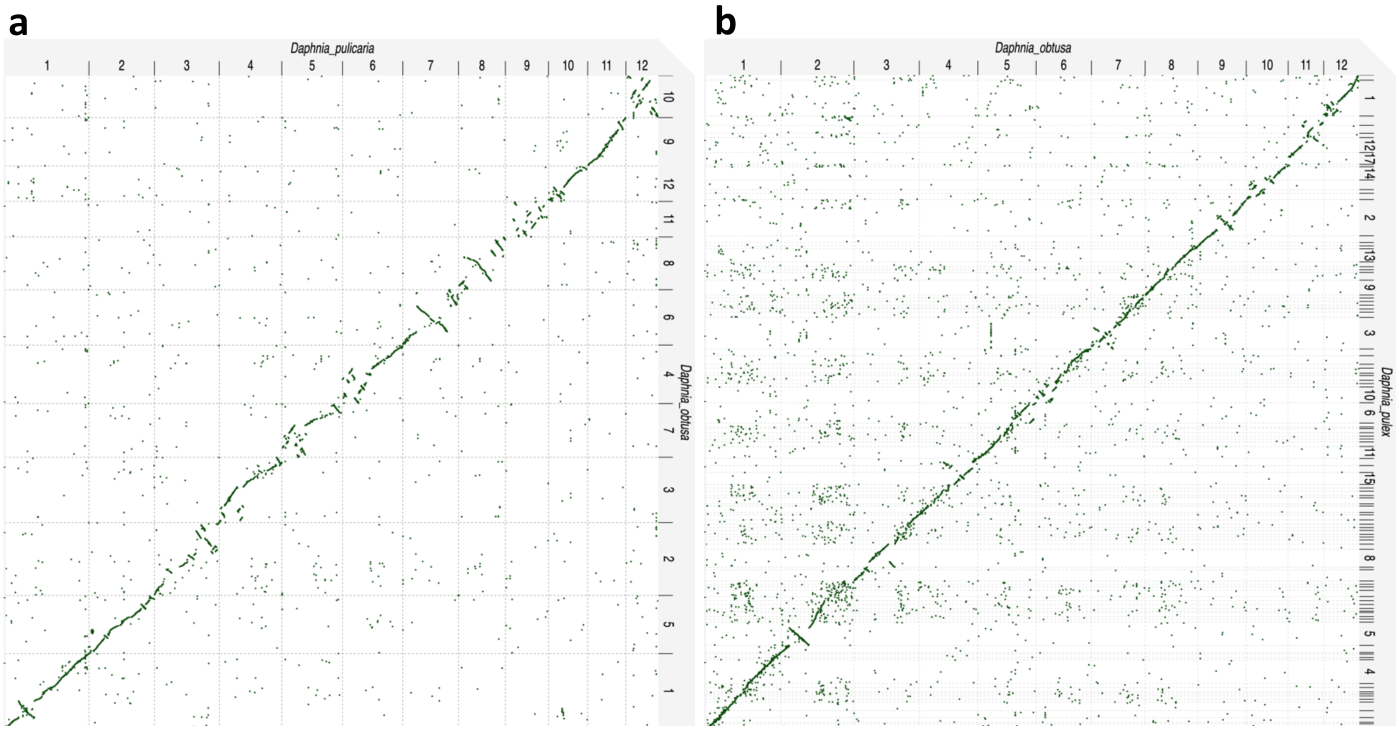


**Supplementary Figure 4. Genome synteny of *D. pulicaria* and *D. pulex* with the outgroup species *D. obtusa***

**(a)** Synteny dotplot visualization of *D. pulicaria* chromosomes (x-axis) and *D. obtusa* chromosomes (y-axis), indicating collinear and rearranged/inverted genomic regions between the two species. **(b)** Synteny dotplot visualization of *D. obtusa* chromosomes (x-axis) and *D. pulex* scaffolds (y-axis), indicating collinear and rearranged/inverted genomic regions between the two species.
